# Anthropocene refugia: integrating history and predictive modelling to assess the space available for biodiversity in a human-dominated world

**DOI:** 10.1101/722132

**Authors:** Sophie Monsarrat, Scott Jarvie, Jens-Christian Svenning

## Abstract

During periods of strong environmental change, some areas may serve as refugia, where components of biodiversity can find protection, persist and potentially expand from should conditions again become favourable. The refugia concept has previously been used in the context of climatic change, to describe climatically stable areas in which taxa survived past Quaternary glacial-interglacial oscillations, or where they might persist in the future under anthropogenic climate change. However, with the recognition that Earth has entered the Anthropocene, an era in which human activities are the dominant driving force on ecosystems, it is critical to also consider human pressures on the environment as factors limiting species distributions. Here, we present a novel concept, Anthropocene refugia, to refer to areas that provide spatial and temporal protection from human activities and that will remain suitable for a given taxonomic unit in the long-term. It integrates a deep-time perspective on species biogeography that provides information on the natural rather than current-day relictual distribution of species, with spatial information on modern and future anthropogenic threats. We define the concept and propose a methodology to effectively identify and map realised and potential current and future refugia, using examples for two megafauna species as a proof of concept. We argue that identifying Anthropocene refugia will improve biodiversity conservation and restoration by allowing better prediction of key areas for conservation and potential for re-expansions today and in the future. More generally, it forms a new conceptual framework to assess and manage the impact of anthropogenic activities on past, current and future patterns of species distributions.

## 2. Introduction

While species distribution patterns have strong signatures tied to natural biotic and abiotic processes, it is becoming increasingly difficult to ignore the role that humans play in shaping the composition of species assemblages across landscapes. With 95% of land having at least some degree of modification by human activities [1], the extent of wilderness areas have declined dramatically, and so have the opportunities to protect and conserve viable populations of many species in their natural habitat. Meanwhile, anthropogenic climate change is predicted to cause range shifts, range contractions and changes in elevational distributions in many organisms [2,3], challenging our approach to biodiversity conservation [4]. Human impacts on the global environment have become so pervasive that a new geological epoch has been proposed: the Anthropocene [5]. Identifying the space available for biodiversity protection and recovery in this human-dominated world is a challenge that requires a comprehensive understanding of the interactions between species’ natural biogeographic patterns and the spatial distribution of anthropogenic pressures.

The terms ‘refuge’ or ‘refugia’ are commonly used to refer to areas where components of biodiversity retreat to, and persist in, under increasing environmental stress, with the potential to re-expand once the stress decreases [6,7]. These concepts have previously been applied in the context of past or contemporary climatic change, to identify areas that are relatively buffered from climatic changes and where components of biodiversity have persisted in the past or may be able to persist in the future. Such areas can act as sources of recolonization when environmental conditions improve, often have long-lasting imprints on species distributions [8], and have therefore become a central focus in much biogeographic research [6]. However, previous uses of the concept have not accounted for the impact of anthropogenic pressures, other than climate change, on species’ biogeographic patterns [6]. With the recognition that human activities are now a major force driving global ecosystems [9], there is a need for incorporating a wider range of human pressures, beyond anthropogenic climate change, as drivers limiting the persistence of species in human-dominated landscapes.

Here, we build on previous uses of the term to introduce a novel concept - Anthropocene refugia - which refers to areas allowing the long-term survival and persistence of organisms that are sensitive to human activities and providing sources for broader recovery if pressures are decreased. It intersects knowledge on the potential distribution of the organism of interest, incorporating prehistoric and historic data on a taxon’s past distribution, with spatially explicit information on current and future anthropogenic pressures. The outcome is the identification of areas where an organism can persist, or be restored to, in the Anthropocene, given its ecology, vulnerability to human activities and the predicted changes in suitability of these areas. This concept will contribute to a much-needed development of more proactive approaches to nature conservation to overcome ongoing species and ecosystem declines and long-term reductions in biodiversity [10,11]. We define this novel conceptual framework and suggest a methodology to identify Anthropocene refugia in practice, identifying the sources of information and available material that can be used for this exercise. We also discuss the applications for species’ conservation, management and restoration, using examples for two megafauna species as case studies and proof of concept.

## 3. Anthropocene refugia: a conceptual framework

The concept of refugia was originally used to study the response of organisms to past periods of glacial-interglacial oscillations of the late Quaternary [6], allowing a greater understanding of observed patterns of species’ biogeography, evolution and demography [e.g. 12–14]. More recently, the term climatic refugia has been applied to contemporary landscapes, referring to locations projected to harbour remnants of present-day climates, which may serve as safe havens for biodiversity under future climate change [7]. In conservation planning, identifying these climate change refugia can help prioritise management efforts in the face of contemporary climate change [15,16].

These definitions, however, only consider climate as the driver of change in species distribution and abundance, both in evolutionary and ecological timeframes [16]. While changes in species’ biogeography during the Quaternary were largely driven by glacial-interglacial oscillations [17], the more recent use of climate change refugia fails to incorporate anthropogenic pressures that, along with climate change, can affect the distribution of species in an increasingly human-dominated world. The concept of Anthropocene refugia overcomes this limitation by incorporating climate change and a wide range of anthropogenic pressures into the identification of refugia. It designates a spatial entity that answers to two qualities: being ecologically suitable for the biodiversity unit considered and having relatively low levels of observed and predicted human pressure to allow its long-term (through several generations) persistence in this area (Figure 1). We differentiate realised and potential refugia, based on whether the refugia are within the taxon’s current range, or within the area where it could potentially persist in the future.

**Figure 1.**
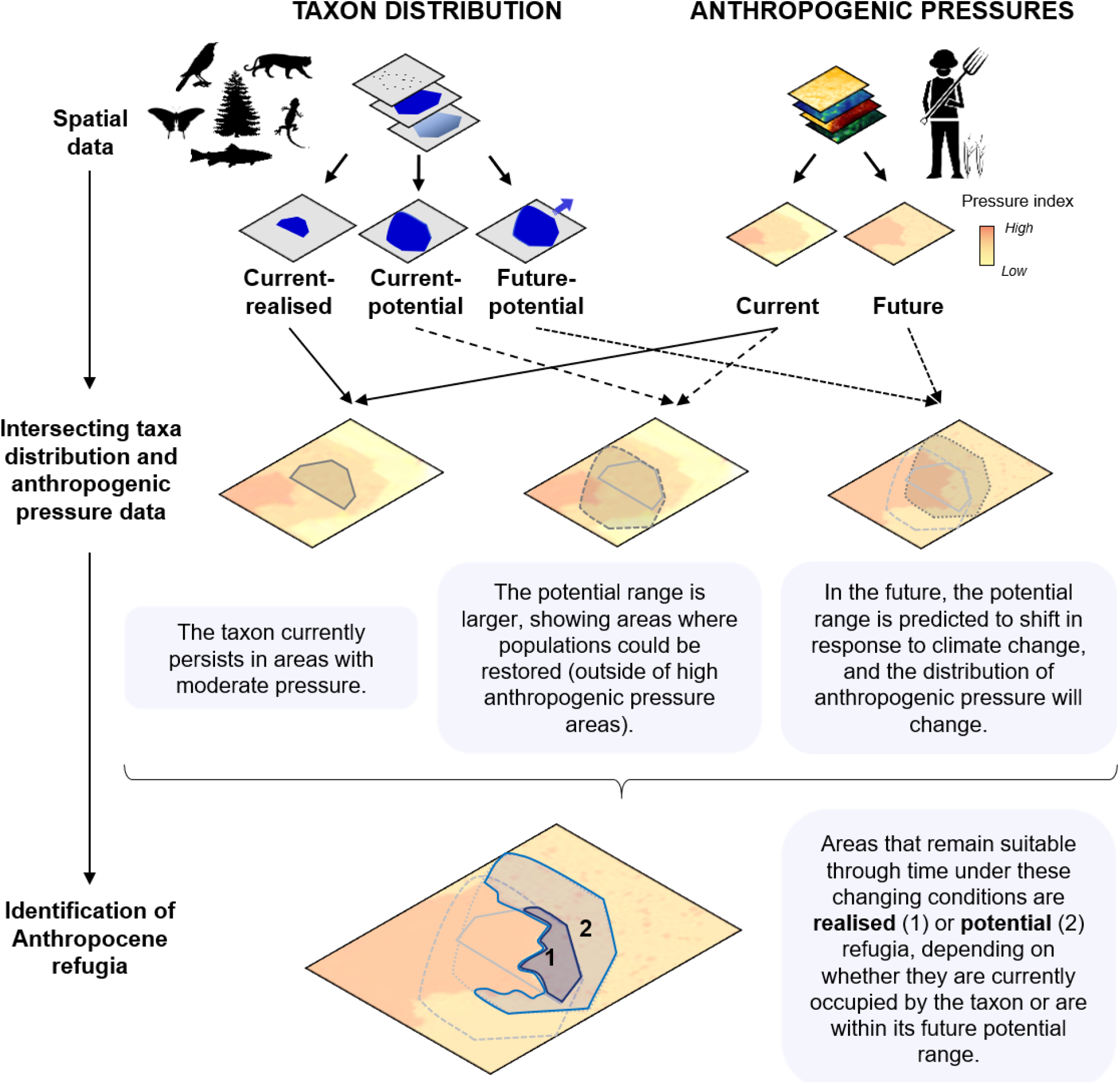
Schematic representation of the Anthropocene refugia concept. Spatially-explicit data on taxon distribution (point data, range maps or predicted distributions) and anthropogenic pressures are combined to map the distribution of realised and potential Anthropocene refugia.

Anthropocene refugia are different from “remnants”, patches of suitable habitat for species intolerant to a human-modified landscape [18], or “coolspots”, areas across species distributions that are free from anthropogenic threats that the species is sensitive to [19], which are both snapshots in time of where species can persist within their current range. In the Anthropocene refugia concept, a strong emphasis is made on identifying the potential range of species, taking into account the natural rather than current-day relictual niche of the species, and the stability of suitable habitat through time. Niche truncation caused by past local extirpations biases our understanding of natural habitat-species relationships. By underestimating the extent of potential suitable habitat, this can affect our understanding of areas available for conservation today [20] and provide unreliable forecasts of distribution changes under future climate change [21]. This case of shifted baseline can be avoided by incorporating prehistoric and historic data on the taxon’s occurrence into estimates of potential distribution. As such, this concept bridges perspectives on taxa’s long-term biogeography with current and future predictions of distribution patterns in human-dominated landscapes.

Anthropocene refugia can be considered both conceptually and as a tool for conservation and management. The Anthropocene refugia concept allows higher-level thinking about the implication and consequences of human activities on species distribution and about the space available for biodiversity in the Anthropocene. Similar to the concept of glacial-interglacial refugia, which allows for a critical examination of current biogeographic patterns with a historical perspective, the identification of Anthropocene refugia will equip the next generations of ecologists with a tool to understand how biogeographic patterns emerge in a human-dominated world. It combines the protection perspective with the restoration (reintroduction or rewilding) perspective, in contrast to more classic conservation approaches which focus on the former [22].

As a tool, Anthropocene refugia can be used in categorisation, decision making, and inference for conservation planning. The approach is complementary to previous efforts to identify priority regions for establishing protected areas or remaining wilderness areas [1,23], but acknowledges that species persistence does not only rely on the existence of formally protected or pressure-free areas [24]. Although protected areas are most certainly a critical component of large-scale conservation planning efforts, some species have demonstrated their ability to tolerate substantial levels of human pressures and survive outside of formally protected areas [25,26]. By enabling persistence, acting as wildlife corridors and serving as a source for future recolonization, refugia are critical to support biodiversity protection in a changing world. Compared to previous efforts to map human footprint [27] or degree of human modification [1], which provide information on the current distribution of human pressures, Anthropocene refugia can incorporate a finer taxonomic resolution, by intersecting these types of information with distribution data for a given taxon. Because individual taxonomic units – populations or species – are often regarded as the fundamental unit of conservation, this makes it a valuable new tool to inform conservation management.

## 4. Identifying Anthropocene refugia

In practice, mapping Anthropocene refugia requires geographic information on the long-term distribution of the taxa of interest and of the relevant anthropogenic pressures (Figure 1). We provide suggestions below for the type of data that could be included and identify some online and open-access datasets from which this information can be retrieved (Supplementary Information).

### Taxon distribution data

Distribution of taxa can be mapped from 1) point data, the locations where a species has been recorded, 2) geographic ranges, the geographic boundaries of the area where a species is known to occur [28] and 3) predicted distributions, the areas where a species is likely to be present as inferred from the suitability of environmental conditions [29]. We refer the reader to existing reviews to understand the attributes, strength and limitations of these different data types in the context of conservation [29–33].

To be useful in mapping Anthropocene refugia, point data need to be transformed into areas, e.g. by intersecting them with geographic units (grid squares or administrative boundaries) or by converting them into geographic ranges (extent of occurrence EOO or area of occupancy AOO [31]), using interpolation and expert knowledge. Predicted distributions that relate species occurrence with environmental conditions based on mechanistic (process-based) or correlative (statistical) niche modelling enable the extrapolation of incomplete point locality data, and the interpolation of habitat suitability measures throughout the range [34,35]. They can provide a more realistic outcome by minimising both commission and omission errors (erroneous indication that a species is present or absent, respectively), but require following high standard practices regarding the choice of species distribution data or mechanistic links describing species’ niche, environmental predictors, and the building and evaluation of the model [36]. Mechanistic and correlative niche models can also be used to predict changes in taxon distribution under future scenarios of climate change [37,38]. From these, hypotheses about the future extent of occurrence of organisms can be derived to inform conservation interventions [e.g. 39], and serve as a basis for mapping Anthropocene refugia.

Niche-truncation caused by past or current anthropogenic impacts may bias our understanding of species’ biogeography and ecological requirements [20,21]. This in turn can lead to underestimating the potential distribution of species and the area available for their conservation [20,40] and bias forecasts of species distribution under future climate change [21]. The distribution data used in the mapping should be informative of the natural rather than current-day relictual range of species, to counter the undesirable effects of the shifting baseline syndrome [41]. This may require a deeper investigation of past local extinctions and truncations in species-environment relationship, which may be informed by integrating knowledge or data on the historical distribution of the species [42].

Long-term biodiversity data, i.e. prehistorical and historical occurrence records collected from archaeological, museum, written and oral sources, can fill gaps in knowledge and inform the natural distribution of organisms, i.e. the range they could occupy today in the absence of human impacts. Fossil records of the late Quaternary can provide information on biodiversity changes as a result of climatic changes and anthropogenic impacts over the last millennia, while historical descriptions and museum specimens collected by literate travellers and naturalists are informative of the more recent past, particularly of the period following European expansion. Finally, traditional ecological knowledge accumulated through many generations of close interactions between people and the natural world can provide local information on the state of ecosystems from the relatively recent past [43]. Ultimately, the temporal cut-off for long-term biodiversity data should be taxon-specific and reflect the timeline and geography of known or suspected impacts as well as the objectives of the study.

If past climate data are available, a multi-temporal calibration approach to identify a taxon’s potential niche from past and present occurrence records can be used to project the range in current environmental conditions using niche modelling [e.g. 39,56], assuming that the niche has remained stable through time [45]. For data-poor regions or taxa, expert knowledge can also bring a useful approximation to the potential natural range of the organism. Finally, a combination of different approaches can prove the most efficient to infer the biogeographic history of species and deduce their current potential range [46,47].

In general, the use of long-term data in ecological analyses is increasing, with positive outcomes to improve our understanding of natural species-habitat relationships and reduce the pervasive effect of the shifting baseline syndrome. As an example, Lentini et al. reconstructed the historical distribution of kākāpō S*trigops habroptilus* using a combination of Holocene fossil records, mid-Holocene climate data and species distribution modelling, identifying areas suitable for reintroductions for this Critically Endangered bird species [44]. Laliberte and Ripple used historical data to identify patterns of range contraction and expansions in 43 species of North American carnivores and ungulates, and showed reduced persistence of species in areas of higher human influence following Euro-American settlement [48]. These types of studies can form the basis of Anthropocene refugia mapping for the taxa they are considering.

All data types are prone to error, incompleteness and biases (temporal, geographical, environmental and taxonomic), which can affect interpretation of distribution patterns. Biases in the Holocene fossil record are examined by Crees et al. in this special issue [49] and discussion on the biases and errors in data extracted from the historical literature and museum specimens can be found in [50–52]. Historical data, while powerful in their potential to highlight neglected aspects of species’ biogeography, require a rigorous methodology for their collection and interpretation to avoid the common pitfalls in historical ecology [53]. Before including long-term data in analyses of species distribution, sources of errors and biases should be thoroughly investigated and corrected for, and the remaining uncertainties acknowledged.

We list examples of online repositories for global species distribution data, including archaeological records, in Supplementary Information. Each of these data sources need to be critically evaluated for their relevance to answer biological questions, according to criteria such as resolution, scale or representativeness of natural range.

### Anthropogenic pressures

The anthropogenic pressures to include in the mapping are those that directly or indirectly have a negative impact on the persistence of the organism of interest and will vary according to the taxon considered and the spatial scale of the study. For example, while land mammals are mostly affected by habitat loss, degradation and harvesting [54], reptiles and amphibians appear to be primarily affected by agriculture and biological resource use, urban development, natural system modification, invasive species and infectious diseases [55,56], and megafaunal species across taxonomic groups are mainly threatened by direct harvesting [57]. Salafsky et al. [58] proposed 11 categories of threats in a unified classification that is being used by the International Union for the Conservation of Nature (IUCN) Red List [59]. It can prove a valuable resource, along with a literature review or expert knowledge, to identify the set of threats that is relevant to the organism of interest.

Recent studies have also used primary data sources on human stressors to build indices of global cumulative impact of human activities on the environment. The Human Footprint project uses for example data on 1) built environments, 2) population density, 3) electric infrastructure, 4) crop lands, 5) pasture lands, 6) roads, 7) railways, and 8) navigable waterways to map the direct and indirect human pressures on the environment globally in 1993 and 2009 [27]. In a recent study, Kennedy et al. [1] provide a cumulative measure of human modification of terrestrial lands based on 13 anthropogenic stressors classified in five major categories: 1) human settlement, 2) agriculture, 3) transportation, 4) mining and energy production, and 5) electrical infrastructure, for a median year of 2016. Spatially explicit global datasets for these two projects are available online [60,61] and are relevant resources that can be used for mapping Anthropocene refugia.

With the growing development of remote-sensing technologies [68], global environment modelling and systematic surveys, spatially-explicit information on current anthropogenic pressures is increasingly made available through publicly available online repositories. We provide a non-exhaustive list of these spatial datasets for different types of human pressures in the Supplementary Information, which can serve as a basis for more case-specific listing of potential data sources. If spatial information for some of the identified threats is unavailable, proxy variables that correlate with the variable of interest can be considered for replacement. The methodology can also be revisited to include additional pressures as more data become available in the future.

New and emerging threats from wildlife trade, changing land use patterns, and the increase in global human populations, may increase pressure on ecosystems in the future. On the other hand, the creation of new protected areas and farmland abandonment may provide additional space for biodiversity [62,63]. In order to identify which areas will allow the long-term persistence of a given organism, these future dynamics need to be incorporated, as much as possible, into the mapping of Anthropocene refugia. Spatially-explicit forecasts of future human population [64], urban expansion [65], global habitat conversion [66] and deforestation [67] are readily available and can be used as a first approximation to understand how the distribution of anthropogenic pressures will change in the future. These datasets however summarise future risks at coarse scales and for different points in time (e.g. from a decade in the future for deforestation risk to a hundred years for human population density), making it a challenge to reconcile them into a single map and to use them as a basis for decision-making at the local scale. Socio-economic, political and ecological trajectories as well as the interactions and retroactive feedbacks between them are generally challenging to model, hindering our capacity to obtain robust predictions of the future distribution of anthropogenic pressures. Given the importance of such predictions for implementing effective conservation and policy measures, the development of methods to produce robust and spatially explicit predictions of the distribution of human activities under different scenarios and over relevant timeframes is an important focus for future research. Release of these predictions in open access format will be critical to improve our ability to identify Anthropocene refugia.

### Mapping Anthropocene refugia

Assigning relative pressure scores to each human pressure variable and intersecting these with species distribution data will result in a taxon-specific map of anthropogenic pressure intensity within the range of the species. Anthropocene refugia lie at the intersection of areas that remains suitable for the organism through time and areas with low levels of observed and predicted anthropogenic pressure. A distinction is made between “realised” refugia that are currently occupied by the organism of interest and predicted to remain suitable in the future, and “potential” refugia, i.e. areas where the taxon is not currently extant but that are within its potential future range under scenarios of future climate change, future human-driven landscape change, and other predicted anthropogenic pressures.

Using thresholds to convert maps of continuous value into binary maps highlighting the specific areas that could act as Anthropocene refugia may be useful for management and decision-making purposes. This approach however leads to loss of valuable information and should be used with caution. If necessary, an option is to calibrate the threshold based on the level of threat that the organism is currently able to sustain within its range [69], the rationale being that an organism will possibly be able to persist in the long-term within areas of lower or similar level of anthropogenic pressure than what it is currently experiencing, with the assumption that the tolerance of humans for its presence will remain similar. This is a conservative approach as refugia could be underestimated for some species that are actually able to sustain higher levels of anthropogenic pressures than what is currently observed. To reflect uncertainty and provide a better decision tool, we recommend testing and reporting variability from using various thresholds. A lower limit on the size of contiguous areas that qualify as refugia for each taxon can also be applied, to meet the requirements for these areas to sustain viable populations [70, but see 71].

In Box 1, we illustrate the Anthropocene refugia concept by mapping realised and potential Anthropocene refugia for two megafauna species, the American bison *Bison bison* and tiger *Panthera tigris*. Both species have undergone massive declines in range and abundance in the past and their persistence is dependent on ongoing conservation programs. Their major ecological roles also make then good candidates for trophic rewilding [72]. While these maps primarily have an illustrative purpose, we briefly discuss potential implications in terms of conservation strategies in Box 1. Details on the methodology to produce these maps are available in the Supplementary Information.

## 5. Applications in conservation

The concept of Anthropocene refugia fills a gap left open by other conservation approaches that only identify climate change refugia or remnant populations to prioritize conservation efforts. The approach is conceptually similar to that proposed by Allan et al. in a recent study, which combined cumulative human impact with current distributions of terrestrial vertebrates to identify hotspots of impacted species richness and coolspots of unimpacted species richness [19]. However, instead of focusing on species’ current range of occurrence, our approach offers a complementary framework in which information from the past is used to understand the potential distribution of a given taxon in the future. We are thus offering a tool to not only define threat mitigation strategies, but to also more fully identify restoration options under future global change and changing anthropogenic pressures. This framework is coherent with the recent designation of 2021–2030 as the “decade of ecosystem restoration” by the United Nations General Assembly and the increasing emphasis put on rewilding as a restoration tool [73]. Identifying those refugia is arguably a challenging exercise and the definition itself is open to further discussion and refinement. With this in mind, we believe there is a high potential for Anthropocene refugia to inform contemporary conservation and restoration, and we suggest possible applications below.

### Realised refugia

Conservation management and restoration within realised Anthropocene refugia is important to maintain existing populations in areas that are predicted to remain suitable for the taxon in the long-term and to promote self-managing, biodiverse ecosystems. This is likely to be facilitated by the lower need for human intervention in areas where taxa are already extant and where the level of anthropogenic pressure is not expected to increase beyond its tolerance level in the near future. This can help identify priority regions for establishing new protected areas, and inform strategies to directly mitigate the threats driving species’ declines [19]. It can feed into decision making for conservation management and conservation planning, e.g. following a similar framework as those proposed for climate change refugia [15,16]. This can also be used to identify candidate sites for reinforcements, i.e. the release of an organism into an existing population of conspecifics to enhance population viability [74]. Nonetheless, in a context of fragmented distributions, there is a possibility that populations within these refugia represent sink populations that would become extinct if they are no longer accessible to dispersers [75], or refugee populations confined to suboptimal habitats, with consequences of decreased fitness and density [76]. The viability of populations in these refuge areas could be assessed using information from paleo- and historical ecology as well as field studies [e.g. 77]. Identifying corridors between refugia and maintaining connectivity between these areas is thus important to allow natural dispersal and improve population persistence [4]. These areas can also be affected by ecological imbalances (e.g. from the loss of ecological interactions [78]), causing a need to restore their functionality, e.g. through trophic rewilding [72]. From a socio-ecological perspective, the implications for local populations of focusing conservation actions on these areas should also be considered [79].

### Potential refugia

Mapping potential Anthropocene refugia can form the basis of a more in-depth evaluation of candidate sites for reintroduction and introduction efforts. For many threatened species, conservation success will rely on the expansion of current relict distributions through reintroductions, a strategy that can be guided by the mapping of Anthropocene refugia. By selecting sites that match the biotic and abiotic needs of the focal species in the long-term, this can also inform restoration strategies such as trophic rewilding, an approach promoting self-regulating ecosystems through the introduction of species to restore top-down trophic interactions and associated trophic cascades [72]. It also opens the possibility to change our approach to conservation interventions and translocations to allow the emergence of novel ecosystems that will be robust to future habitat shifts and changing anthropogenic impacts [80]. This may involve planning corridors to allow colonisation through natural dispersal or implementing measures to introduce species outside of their historical range, an approach called assisted colonisation [81].

Identifying areas that are most likely to be recolonised in the near future is also important to anticipate the complex impacts this could have on ecosystems as well as interactions with society. The socio-ecological implications of some species’ recolonising part of their historical range or occupying new areas should be carefully considered, in particular for species often involved in human-wildlife conflicts, such as large carnivores [82]. Unified socio-ecological approaches that explicitly acknowledge competing perspectives between wildlife conservation and social and governance contexts can be applied to provide new insights into management options for sustainable human–wildlife coexistence [83].

## 6. Challenges and opportunities in mapping Anthropocene refugia

The highly dynamic state that characterizes the Anthropocene hinders our ability to make predictions for the direction and intensity of changes in the global environment beyond the next few decades. Future global warming, with temperatures predicted to exceed Quaternary levels, will have potential knock-on effects on the entire biosphere and unpredictable consequences for the survival of individual species [84]. This challenges our ability to predict the location of stable Anthropocene refugia, a significant limitation that will not easily be resolved. In this context of uncertainty, focusing conservation efforts on areas that are likely to remain suitable in the near future might be our best option to increase the chances of survival for sensitive taxa and maintain a genetic diversity that will allow future recovery. It also gains time for conservationists and managers to develop long-term solutions for the survival of populations and species. The historical pressures that caused a taxon’s local extinction must no longer exist locally for an area to be considered a suitable candidate for restoration. As some threats can be difficult to explicitly include in the mapping exercise, a thorough investigation of past and existing impacts from human activities is necessary before actual conservation recommendations can be proposed. Opportunities to improve the mapping of long-term refugia will increase as we gain a better understanding of taxa distributions’ response to climate change and with the future release of spatial data on predicted changes in anthropogenic pressures. As these are integrated into the analysis, one can move from a paradigm focused on places to one focused on dynamics.

Another important consideration is the scale at which Anthropocene refugia should be identified and managed. The geopolitical scale used in conservation has an enormous influence on the identification of management areas, with for example low congruence between the network of conservation areas identified at the broad regional *vs* the fine local scale [85]. In practice, the definition of conservation targets is often site-specific. For example, the Natura 2000 network of protected areas, one of the pillars of the European Community conservation policy, emphasizes the management of specific sites [86]. Finer-scale refugia maps based on local-scale and high-resolution distribution information could thus prove relevant for the identification of land units that will form the basis of management decisions in this context. However, this site-specific approach does not always provide an efficient prioritisation in relation to broader biodiversity concerns [87,88]. Smaller-scale analyses may lead to suboptimal prioritisation with respect to the value for global biodiversity and cost effectiveness, calling for a wider historical and geographical context to contextualise management measures. This demonstrates how conservation strategies can be identified over a wide range of scales and how identifying Anthropocene refugia at both a fine and broad resolution can prove useful for conservation. Ultimately, decisions relative to scale will depend on the objectives of each project and the availability of relevant datasets. The rationale for these choices needs to be explicit and argued, in order to withstand scrutiny and allow for future refinement.

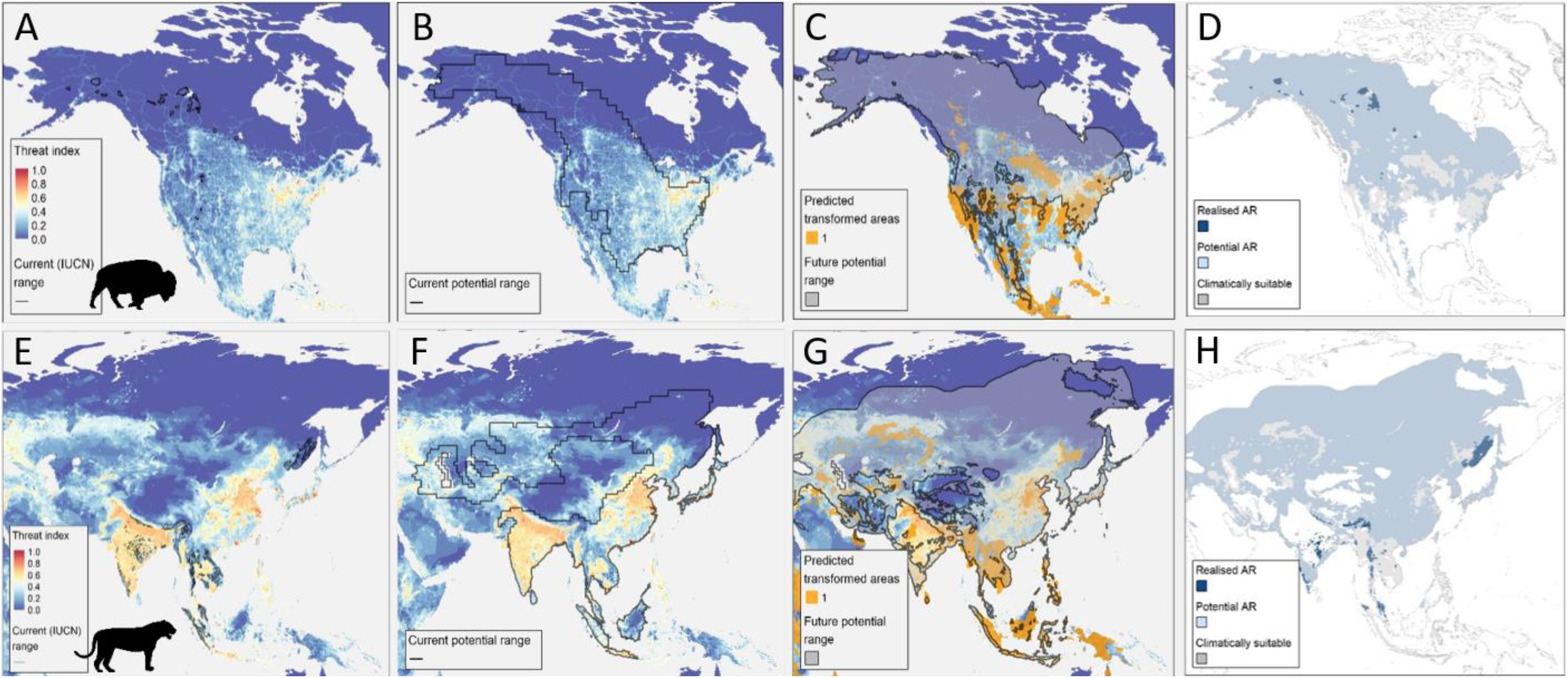

#### Box 1.

##### Illustration of Anthropocene refugia mapping for the American bison *Bison bison* (A to D) and Tiger *Panthera tigris* (E to H)

Maps A-C and E-G are based on the taxa-specific index of current anthropogenic threat, on which we overlaid the areas with a high probability of being converted into cropland or built areas in the future (predicted future transformed areas) for maps C and G. A and E show the current (IUCN) ranges where each species is extant today, B and F show the current potential distributions, and C and G show the potential future range under climate change. D and H show the locations of realised and potential Anthropocene refugia (AR) for the two species, calculated by applying a threshold on the threat index within the potential future range of the species equal to the 95% percentile of the threat level currently experienced by the species. Grey areas are predicted to be climatically suitable for the species in the future but have a value of threat index above the reported threshold. See Supplementary Information for details about the methodology and maps of AR using different thresholds.

##### American bison

The American bison currently persists in very fragmented and highly managed populations in the United States and Canada. The areas within the northernmost part of the range, which are within the future predicted range for the species and with low levels of threats now and in the future, are realised refugia for the species. Conservation actions could focus on managing and reinforcing populations in these areas, e.g. by enforcing protection and building corridors between these populations. The climatically suitable area is also predicted to expand in Canada and Alaska under future climate change, and land in these areas is not predicted to be transformed in the near future, making these regions potential refugia. These are good candidate areas to restore bison populations through reintroductions or assisted colonisation. On the contrary, the southern part of the current potential range has a high level of threats and the range of the species is predicted to contract and fragment under future climate change. Conservation and restoration in this area will thus require greater efforts.

##### Tiger

The current range of the tiger is very fragmented and located in areas surrounded with high anthropogenic pressures (e.g. in India). Some of these areas represent realised refugia because they are predicted to remain within the range of the species under future climate change and the anthropogenic pressures are not expected to increase (e.g. protected areas in India, Bhutan, China and southeast Asia and in the Russian Far East). Threat levels in northeast China, Russia, Mongolia and Kazakhstan are low and the climate suitability is predicted to remain stable. These are potential candidate areas for restoration, although a more thorough investigation of historical threats that drove the species’ local extinction in these areas would be needed to shed light on their current suitability. Other areas are part of the current potential range of the species, but may become unsuitable in the future because of predicted range shifts under climate change (e.g. Borneo) or a predicted increase in land-conversion (Southeast Asia and part of Indonesia). These elements could be taken into account for establishing conservation priorities to ensure the species' long-term persistence.

## 7. Conclusion

We here introduce a novel concept – Anthropocene refugia – to account for the role of anthropogenic pressures in defining realised and potential refugia for biodiversity in a human-dominated planet. Our main intention is to call for a better consideration of the full range of anthropogenic pressures, beyond climate change, to identify refugia in the Anthropocene, while at the same time assessing re-expansion possibilities. We also emphasise the importance of considering a long-term perspective in defining a taxon’s potential distribution, to overcome the shifting baseline syndrome that affects our understanding of natural biogeographical patterns, limiting many assessments of restoration options. We highlight key possible applications of the concept with special emphasis on its potential to inform restoration approaches such as trophic rewilding, as well as nature management more generally, in the face of contemporary climate change. It is our hope to see a larger range of potential applications of this approach discussed and implemented in the future. Despite the many challenges, we argue that the Anthropocene refugia concept has the potential to bridge important gaps in our perspectives on the past (the distribution and ecology of wildlife prior to severe human pressure), present (where wildlife occur now in a human-dominated world) and future (where society could hopefully allow wildlife to exist in a reconciling world), thus representing both an important concept to reflect on our coexistence with wildlife on this planet, and an integral component of the conservation and restoration toolbox to protect and promote biodiversity in the Anthropocene.

## Acknowledgments

This work is a contribution to the Carlsberg Foundation Semper Ardens project MegaPast2Future (grant CF16-0005 to JCS) and to the VILLUM Investigator project “Biodiversity Dynamics in a Changing World” funded by VILLUM FONDEN (grant 16549 to JCS). We thank Emilio Berti for thoughts on the species distribution modelling and MegaPast2Future project members for insightful comments.

## References

1. Kennedy CM, Oakleaf JR, Theobald DM, Baruch-Mordo S, Kiesecker J. 2019 Managing the middle: A shift in conservation priorities based on the global human modification gradient. Glob. Change Biol. (doi:10.1111/gcb.14549)

2. Freeman BG, Lee-Yaw JA, Sunday JM, Hargreaves AL. 2018 Expanding, shifting and shrinking: The impact of global warming on species’ elevational distributions. Glob. Ecol. Biogeogr. (doi:10.1111/geb.12774)

3. Lenoir J, Svenning J-C. 2015 Climate-related range shifts - a global multidimensional synthesis and new research directions. Ecography 38, 15–28. (doi:10.1111/ecog.00967)

4. Heller NE, Zavaleta ES. 2009 Biodiversity management in the face of climate change: A review of 22 years of recommendations. Biol. Conserv. 142, 14–32. (doi:10.1016/j.biocon.2008.10.006)

5. Crutzen PJ. 2002 Geology of mankind. Nature 415, 27.

6. Bennett KD, Provan J. 2008 What do we mean by ‘refugia’? Ice Age Refug. Quat. Extinctions Issue Quat. Evol. Palaeoecol. 27, 2449–2455. (doi:10.1016/j.quascirev.2008.08.019)

7. Keppel G, Van Niel KP, Wardell-Johnson GW, Yates CJ, Byrne M, Mucina L, Schut AGT, Hopper SD, Franklin SE. 2012 Refugia: identifying and understanding safe havens for biodiversity under climate change. Glob. Ecol. Biogeogr. 21, 393–404. (doi:10.1111/j.1466-8238.2011.00686.x)

8. Graham CH, Moritz C, Williams SE. 2006 Habitat history improves prediction of biodiversity in rainforest fauna. Proc. Natl. Acad. Sci. 103, 632–636. (doi:10.1073/pnas.0505754103)

9. Vitousek PM, Mooney HA, Lubchenco J, Melillo JM. 1997 Human Domination of Earth’s Ecosystems. Science 277, 494–499. (doi:10.1126/science.277.5325.494)

10. Davis M, Faurby S, Svenning J-C. 2018 Mammal diversity will take millions of years to recover from the current biodiversity crisis. Proc. Natl. Acad. Sci. 115, 11262–11267. (doi:10.1073/pnas.1804906115)

11. Svenning J-C. 2018 Proactive conservation and restoration of botanical diversity in the Anthropocene’s “rambunctious garden”. Am. J. Bot. 105, 963–966. (doi:10.1002/ajb2.1117)

12. Svenning J-C, Normand S, Kageyama M. 2008 Glacial refugia of temperate trees in Europe: insights from species distribution modelling. J. Ecol. 96, 1117–1127. (doi:10.1111/j.1365-2745.2008.01422.x)

13. Hewitt GM. 2004 Genetic consequences of climatic oscillations in the Quaternary. Philos. Trans. R. Soc. Lond. B. Biol. Sci. 359, 183–195. (doi:10.1098/rstb.2003.1388)

14. Wallis GP, Waters JM, Upton P, Craw D. 2016 Transverse Alpine Speciation Driven by Glaciation. Trends Ecol. Evol. 31, 916–926. (doi:10.1016/j.tree.2016.08.009)

15. Morelli TL et al. 2016 Managing Climate Change Refugia for Climate Adaptation. PLOS ONE 11, e0159909. (doi:10.1371/journal.pone.0159909)

16. Keppel G, Mokany K, Wardell-Johnson GW, Phillips BL, Welbergen JA, Reside AE. 2015 The capacity of refugia for conservation planning under climate change. Front. Ecol. Environ. 13, 106–112. (doi:10.1890/140055)

17. Stewart JR, Lister AM, Barnes I, Dalén L. 2010 Refugia revisited: individualistic responses of species in space and time. Proc. R. Soc. B Biol. Sci. 277, 661–671. (doi:10.1098/rspb.2009.1272)

18. Keppel G, Wardell-Johnson GW. 2012 Refugia: keys to climate change management. Glob. Change Biol. 18, 2389–2391. (doi:10.1111/j.1365-2486.2012.02729.x)

19. Allan JR, Watson JEM, Marco MD, Possingham HP, Venter O. 2019 Hotspots of human impact on threatened terrestrial vertebrates. PLoS Biol., 18.

20. Monsarrat S, Novellie P, Rushworth I, Kerley GIH. 2019 Shifted distribution baselines: neglecting long-term biodiversity records risks overlooking potentially suitable habitat for conservation management. bioRxiv, 565929. (doi:10.1101/565929)

21. Faurby S, Araújo MB. 2018 Anthropogenic range contractions bias species climate change forecasts. Nat. Clim. Change 8, 252–256. (doi:10.1038/s41558-018-0089-x)

22. Margules CR, Pressey RL. 2000 Systematic conservation planning. Nature 405, 243–253.

23. Allan JR, Venter O, Watson JEM. 2017 Temporally inter-comparable maps of terrestrial wilderness and the Last of the Wild. Sci. Data 4, 170187. (doi:10.1038/sdata.2017.187)

24. Dudley N, Jonas H, Nelson F, Parrish J, Pyhälä A, Stolton S, Watson JEM. 2018 The essential role of other effective area-based conservation measures in achieving big bold conservation targets. Glob. Ecol. Conserv. 15, e00424. (doi:10.1016/j.gecco.2018.e00424)

25. Galán-Acedo C, Arroyo-Rodríguez V, Andresen E, Verde Arregoitia L, Vega E, Peres CA, Ewers RM. 2019 The conservation value of human-modified landscapes for the world’s primates. Nat. Commun. 10. (doi:10.1038/s41467-018-08139-0)

26. López-Bao JV, Kaczensky P, Linnell JDC, Boitani L, Chapron G. 2015 Carnivore coexistence: Wilderness not required. Science 348, 871. (doi:10.1126/science.348.6237.871-b)

27. Venter O et al. 2016 Global terrestrial Human Footprint maps for 1993 and 2009. Sci. Data 3, 160067. (doi:10.1038/sdata.2016.67)

28. Gaston KJ. 1991 How Large Is a Species’ Geographic Range? Oikos 61, 434. (doi:10.2307/3545251)

29. Rondinini C, Wilson KA, Boitani L, Grantham H, Possingham HP. 2006 Tradeoffs of different types of species occurrence data for use in systematic conservation planning: Species data for conservation planning. Ecol. Lett. 9, 1136–1145. (doi:10.1111/j.1461-0248.2006.00970.x)

30. Boitani L, Maiorano L, Baisero D, Falcucci A, Visconti P, Rondinini C. 2011 What spatial data do we need to develop global mammal conservation strategies? Philos. Trans. R. Soc. B Biol. Sci. 366, 2623–2632. (doi:10.1098/rstb.2011.0117)

31. Gaston KJ, Fuller RA. 2009 The sizes of species’ geographic ranges. J. Appl. Ecol. 46, 1–9. (doi:10.1111/j.1365-2664.2008.01596.x)

32. Bombi P, Luiselli L, D’Amen M. 2011 When the method for mapping species matters: defining priority areas for conservation of African freshwater turtles: Prioritization of African turtles. Divers. Distrib. 17, 581–592. (doi:10.1111/j.1472-4642.2011.00769.x)

33. Rodríguez JP, Brotons L, Bustamante J, Seoane J. 2007 The application of predictive modelling of species distribution to biodiversity conservation. Divers. Distrib. 13, 243–251. (doi:10.1111/j.1472-4642.2007.00356.x)

34. Elith J, Leathwick JR. 2009 Species Distribution Models: Ecological Explanation and Prediction Across Space and Time. Annu. Rev. Ecol. Evol. Syst. 40, 677–697. (doi:10.1146/annurev.ecolsys.110308.120159)

35. Kearney M, Porter W. 2009 Mechanistic niche modelling: combining physiological and spatial data to predict species’ ranges. Ecol. Lett. 12, 334–350. (doi:10.1111/j.1461-0248.2008.01277.x)

36. Araújo MB et al. 2019 Standards for distribution models in biodiversity assessments. Sci. Adv. 5, eaat4858. (doi:10.1126/sciadv.aat4858)

37. Pearson RG, Dawson TP. 2003 Predicting the impacts of climate change on the distribution of species: are bioclimate envelope models useful? Glob. Ecol. Biogeogr. 12, 361–371.

38. Kearney MR, Wintle BA, Porter WP. 2010 Correlative and mechanistic models of species distribution provide congruent forecasts under climate change: Congruence of correlative and mechanistic distribution models. Conserv. Lett. 3, 203–213. (doi:10.1111/j.1755-263X.2010.00097.x)

39. Jarvie S, Svenning J-C. 2018 Using species distribution modelling to determine opportunities for trophic rewilding under future scenarios of climate change. Philos. Trans. R. Soc. B Biol. Sci. 373, 10. (doi:http://dx.doi.org/10.1098/rstb.2017.0446)

40. Nüchel J, Bøcher PK, Xiao W, Zhu A-X, Svenning J-C. 2018 Snub-nosed monkeys (Rhinopithecus): potential distribution and its implication for conservation. Biodivers. Conserv. 27, 1517–1538. (doi:10.1007/s10531-018-1507-0)

41. Soga M, Gaston KJ. 2018 Shifting baseline syndrome: causes, consequences, and implications. Front. Ecol. Environ. 16, 263–263.

42. Turvey ST, Crees JJ, Di Fonzo MMI. 2015 Historical data as a baseline for conservation: reconstructing long-term faunal extinction dynamics in Late Imperial–modern China. Proc. R. Soc. B Biol. Sci. 282, 20151299. (doi:10.1098/rspb.2015.1299)

43. Eckert LE, Ban NC, Frid A, McGreer M. 2018 Diving back in time: Extending historical baselines for yelloweye rockfish with Indigenous knowledge. Aquat. Conserv. Mar. Freshw. Ecosyst. 28, 158–166. (doi:10.1002/aqc.2834)

44. Lentini PE, Stirnemann IA, Stojanovic D, Worthy TH, Stein JA. 2018 Using fossil records to inform reintroduction of the kakapo as a refugee species. Biol. Conserv. 217, 157–165. (doi:10.1016/j.biocon.2017.10.027)

45. Nogués-Bravo D. 2009 Predicting the past distribution of species climatic niches. Glob. Ecol. Biogeogr. 18, 521–531. (doi:10.1111/j.1466-8238.2009.00476.x)

46. Faurby S, Svenning J-C. 2015 Historic and prehistoric human-driven extinctions have reshaped global mammal diversity patterns. Divers. Distrib. 21, 1155–1166. (doi:10.1111/ddi.12369)

47. Hoban S, Dawson A, Robinson JD, Smith AB, Strand AE. 2019 Inference of biogeographic history by formally integrating distinct lines of evidence: genetic, environmental niche and fossil. Ecography, ecog.04327. (doi:10.1111/ecog.04327)

48. Laliberte AS, Ripple WJ. 2004 Range Contractions of North American Carnivores and Ungulates. BioScience 54, 123. (doi:10.1641/0006-3568(2004)054[0123:RCONAC]2.0.CO;2)

49. Crees JJ, Collen B, Turvey ST. 2019 Bias, incompleteness, and the “known unknowns” in the Holocene faunal record. Philos. Trans. R. Soc. B Biol. Sci. **This issue**.

50. Monsarrat S, Kerley GIH. 2018 Charismatic species of the past: Biases in reporting of large mammals in historical written sources. Biol. Conserv. 223, 68–75. (doi:10.1016/j.biocon.2018.04.036)

51. Monsarrat S, Boshoff AF, Kerley GIH. 2018 Accessibility maps as a tool to predict sampling bias in historical biodiversity occurrence records. Ecography (doi:10.1111/ecog.03944)

52. Newbold T. 2010 Applications and limitations of museum data for conservation and ecology, with particular attention to species distribution models. Prog. Phys. Geogr. 34, 3–22. (doi:10.1177/0309133309355630)

53. Pooley S. 2018 Descent with modification: Critical use of historical evidence for conservation. Conserv. Lett., e12437. (doi:10.1111/conl.12437)

54. Schipper J et al. 2008 The status of the world’s land and marine mammals: diversity, threat, and knowledge. Science 322, 225–230. (doi:10.1126/science.1165115)

55. Böhm M et al. 2013 The conservation status of the world’s reptiles. Biol. Conserv. 157, 372–385. (doi:10.1016/j.biocon.2012.07.015)

56. Daszak P, Cunningham AA, Hyatt AD. 2003 Infectious disease and amphibian population declines. Divers. Distrib., 10.

57. Ripple WJ et al. 2019 Are we eating the world’s megafauna to extinction? Conserv. Lett., e12627. (doi:10.1111/conl.12627)

58. Salafsky N et al. 2008 A Standard Lexicon for Biodiversity Conservation: Unified Classifications of Threats and Actions: *Classifications of Threats & Actions*. Conserv. Biol. 22, 897–911. (doi:10.1111/j.1523-1739.2008.00937.x)

59. IUCN. 2018 The IUCN Red List of Threatened Species. Version 2018-2. http://www.iucnredlist.org.

60. Venter O et al. 2016 Data from: Global terrestrial Human Footprint maps for 1993 and 2009. (doi:10.5061/dryad.052q5.2)

61. Christina M. Kennedy, James Oakleaf, David M. Theobald, Sharon Baruch-Mordo, Joseph Kiesecker. 2018 Global Human Modification. (doi:10.6084/m9.figshare.7283087.v1)

62. Queiroz C, Beilin R, Folke C, Lindborg R. 2014 Farmland abandonment: threat or opportunity for biodiversity conservation? A global review. Front. Ecol. Environ. 12, 288–296. (doi:10.1890/120348)

63. Ceauşu S, Hofmann M, Navarro LM, Carver S, Verburg PH, Pereira HM. 2015 Mapping opportunities and challenges for rewilding in Europe. Conserv. Biol. 29, 1017–1027. (doi:10.1111/cobi.12533)

64. Jones B, O’Neill BC. 2016 Spatially explicit global population scenarios consistent with the Shared Socioeconomic Pathways. Environ. Res. Lett. 11, 084003. (doi:10.1088/1748-9326/11/8/084003)

65. Seto K, Güneralp B, Hutyra LR. 2015 Global Grid of Probabilities of Urban Expansion to 2030.

66. Oakleaf JR, Kennedy CM, Baruch-Mordo S, West PC, Gerber JS, Jarvis L, Kiesecker J. 2015 A World at Risk: Aggregating Development Trends to Forecast Global Habitat Conversion. PLOS ONE 10, e0138334. (doi:10.1371/journal.pone.0138334)

67. Hewson J, Crema S, González-Roglich M, Tabor K, Harvey C. 2019 New 1 km Resolution Datasets of Global and Regional Risks of Tree Cover Loss. Land 8, 14. (doi:10.3390/land8010014)

68. Kerr JT, Ostrovsky M. 2003 From space to species: ecological applications for remote sensing. Trends Ecol. Evol. 18, 299–305. (doi:10.1016/S0169-5347(03)00071-5)

69. Escobar LE, Awan MN, Qiao H. 2015 Anthropogenic disturbance and habitat loss for the red-listed Asiatic black bear (Ursus thibetanus): Using ecological niche modeling and nighttime light satellite imagery. Biol. Conserv. 191, 400–407. (doi:10.1016/j.biocon.2015.06.040)

70. Pe’er G et al. 2014 Toward better application of minimum area requirements in conservation planning. Biol. Conserv. 170, 92–102. (doi:10.1016/j.biocon.2013.12.011)

71. Wintle BA et al. 2019 Global synthesis of conservation studies reveals the importance of small habitat patches for biodiversity. Proc. Natl. Acad. Sci. 116, 909–914. (doi:10.1073/pnas.1813051115)

72. Svenning J-C et al. 2016 Science for a wilder Anthropocene: Synthesis and future directions for trophic rewilding research. Proc. Natl. Acad. Sci. 113, 898–906.

73. Perino A et al. 2019 Rewilding complex ecosystems. Science 364, eaav5570. (doi:10.1126/science.aav5570)

74. Seddon PJ, Griffiths CJ, Soorae PS, Armstrong DP. 2014 Reversing defaunation: Restoring species in a changing world. Science 345, 406–412. (doi:10.1126/science.1251818)

75. Fahrig L, work(s): GMR. 1994 Conservation of Fragmented Populations. Conserv. Biol. 8, 50–59.

76. Kerley GIH, Kowalczyk R, Cromsigt JPGM. 2012 Conservation implications of the refugee species concept and the European bison: king of the forest or refugee in a marginal habitat? Ecography 35, 519–529. (doi:10.1111/j.1600-0587.2011.07146.x)

77. Han H et al. 2019 Diet Evolution and Habitat Contraction of Giant Pandas via Stable Isotope Analysis. Curr. Biol. 29, 664–669.e2. (doi:10.1016/j.cub.2018.12.051)

78. Valiente-Banuet A et al. 2015 Beyond species loss: the extinction of ecological interactions in a changing world. Funct. Ecol. 29, 299–307. (doi:10.1111/1365-2435.12356)

79. Burgess ND et al. 2007 Correlations among species distributions, human density and human infrastructure across the high biodiversity tropical mountains of Africa. Biol. Conserv. 134, 164–177. (doi:10.1016/j.biocon.2006.08.024)

80. Hobbs RJ, Higgs E, Harris JA. 2009 Novel ecosystems: implications for conservation and restoration. Trends Ecol. Evol. 24, 599–605. (doi:10.1016/j.tree.2009.05.012)

81. IUCN/SSC. 2013 Guidelines for Reintroductions and Other Conservation Translocations., viiii + 57— pp.

82. Bruskotter JT, Shelby LB. 2010 Human Dimensions of Large Carnivore Conservation and Management: Introduction to the Special Issue. Hum. Dimens. Wildl. 15, 311–314. (doi:10.1080/10871209.2010.508068)

83. Ceauşu S, Graves RA, Killion AK, Svenning J-C, Carter NH. 2019 Governing trade-offs in ecosystem services and disservices to achieve human–wildlife coexistence. Conserv. Biol. 33, 543–553. (doi:10.1111/cobi.13241)

84. Moritz C, Agudo R. 2013 The Future of Species Under Climate Change: Resilience or Decline? Science 341, 504–508. (doi:10.1126/science.1237190)

85. Erasmus BFN, Freitag S, Gaston KJ, Erasmus BH, Jaarsveld AS van. 1999 Scale and conservation planning in the real world. Proc. R. Soc. Lond. B Biol. Sci. 266, 315–319. (doi:10.1098/rspb.1999.0640)

86. EC—European Commission. 2000 Managing Natura 2000 sites: the provisions of Article 6 of the ‘Habitats’ Directive 92/43/EEC. Office for Official Publications of the European Communities: Luxembourg.

87. Albuquerque FS, Assunção-Albuquerque MJT, Cayuela L, Zamora R, Benito BM. 2013 European Bird distribution is “well” represented by Special Protected Areas: Mission accomplished? Biol. Conserv. 159, 45–50. (doi:10.1016/j.biocon.2012.10.012)

88. Bladt J, Strange N, Abildtrup J, Svenning J-C, Skov F. 2009 Conservation efficiency of geopolitical coordination in the EU. J. Nat. Conserv. 17, 72–86. (doi:10.1016/j.jnc.2008.12.003)

