## Supplementary Information for "Anthropocene refugia: integrating history and predictive modelling to assess the space available for biodiversity in a human-dominated world" (from another version of this preprint: version 1, 2019-08-01)

**Supplementary Information 1.** Examples of sources for global species distribution datasets available through open-access online repositories.

| Data type | Data sources | Description | Taxonomic group | Ref |
|---|---|---|---|---|
| Point data | GBIF | The Global Biodiversity Information Facility (GBIF) is an international open data infrastructure that provides open access to modern and fossil species occurrence records for all types of life on Earth. With more than one billion occurrence records for almost 6 million species, obtained from 42,339 datasets (as of February 2019), it is the most widely used repository for biodiversity data. | Almost 6 million species, from all taxonomic groups | [1] |
| | The Paleobiology Database (PBDB) | The PBDB provides global, collection-based occurrence and taxonomic data for organisms of all geological ages. | All | https://paleobiodb.org |
| | Neotoma Paleoecology Database | Neotoma is a public database containing fossil data from the Holocene, Pleistocene, and Pliocene, or approximately the last 5.3 million years. It hosts the Global Pollen Database (GPD), the North American Plant Macrofossil Database (NAPMD), and the database of mammalian fauna (FAUNMAP). | Mammals, pollens | https://www.neotomadb.org/ |
| Geographic ranges | IUCN Red List | The International Union for Conservation of Nature (IUCN) Red List assesses species' conservation status by assembling information about range, population size, habitat and ecology, use and/or trade, threats, and conservation actions. The inventory contains global assessments for over 96,500 species, of which about two-thirds have spatial data on geographic ranges (as of February 2019). Though not required by the Red List, some assessments | All birds, mammals, amphibian and reef building corals, and representative samples of fish, invertebrate, reptile, butterfly and plant taxa | [2] |

| | | | | |
|---|---|---|---|---|
| | | provide information on the historical range of species, with areas coded as "Extinct" or "Possibly Extinct". | | |
| | PHYLACINE | PHYLACINE, The Phylogenetic Atlas of Mammal Macroecology, contains phylogenies, range maps, trait data, and threat status for all 5,831 known mammal species that lived since the last interglacial. The database includes current ranges and counterfactual present natural ranges, i.e. estimates of where species would live today in the absence of anthropogenic pressures. | Mammals | [3] |
| Predicted distribution | BIEN | The Botanical Information and Ecology Network (BIEN) brings together data on plant distribution, abundance and traits from specimens, vegetation inventories, and regional checklists as well as derived products such as species range maps. | Plants | [4,5] |
| | AquaMaps | AquaMaps is a tool for generating model-based, large-scale predictions of natural occurrences of species. It provides standardized distribution maps for over 25,000 species of fishes, marine mammals and invertebrates, including maps of native range and habitat suitability. | Marine species | [6] |

**Supplementary Information 2.** Examples of publicly available global spatially-explicit datasets for contemporary human pressures, following the classification of threats in Salafsky [7].

| Threat | Proxy | Dataset (resolution - when applicable) | Link |
|---|---|---|---|
| Residential and commercial development | Artificial surfaces and associated areas (urban areas > 50%) | GlobCover: Global Land Cover Map (300 metres) | http://due.esrin.esa.int/page_globcover.php |
| Agriculture and aquaculture | Cropland | GlobCover: Global Land Cover Map (300 metres) | http://due.esrin.esa.int/page_globcover.php |
| | Cropland | Unified Cropland Layer (250 metres) | https://www.mdpi.com/2072-4292/7/6/7959 |
| | Livestock production system | Global map of livestock distribution and production system (3 arc-minute) | https://livestock.geo-wiki.org/home-2/ |
| Energy production and mining | Energy production features (mining, wind turbines, oil wells, powerlines) | Open Street Map | https://www.openstreetmap.org/ |
| Transportation and service corridors | Road network | Global Roads Inventory Project (GRIP) dataset | https://www.globio.info/download-grip-dataset |
| | Rail network | Open Street Map | https://www.openstreetmap.org/ |
| | Shipping lanes | 2008 shipping lanes | https://knb.ecoinformatics.org/view/doi:10.5063/F19Z92TW |
| Biological resource use | NPP appropriation | Global Patterns in Human Appropriation of Net Primary Productivity (HANPP) (0.25 x 0.25 degrees) | http://sedac.ciesin.columbia.edu/data/set/hanpp-human-appropriation-net-primary-productivity |
| Human intrusions and disturbance | Human population size | WorldPop Project Population Data: Estimated Residential Population per 100 x 100m Grid Square (3 arc seconds) | https://www.worldpop.org/ |
| | Human population density | Gridded Population of the World (GPW), v4 (30 arc-seconds) | http://sedac.ciesin.columbia.edu/data/set/gpw-v4-population-density-rev10 |
| | Accessibility | Travel time to major cities: A global map of Accessibility (30 arc-seconds) | https://forobs.jrc.ec.europa.eu/products/gam/download.php |
| Natural system modification | Dams and water management | Global Reservoir and Dam Database | http://sedac.ciesin.columbia.edu/data/set/grand-v1-dams-rev01 |
| Invasive and other problematic species and genes | Spatial range of invasive species | IUCN range maps | https://www.iucnredlist.org/resources/spatial-data-download |
| Pollution | Air pollution | HTAP_v2.2: a mosaic of regional and global emission grid maps (0.1 x 0.1 degrees) | https://data.europa.eu/euodp/data/dataset/jrc-edgar-htap_v2-2 |
| | Light pollution | World atlas of artificial night sky brightness (30 arc-seconds) | http://advances.sciencemag.org/content/2/6/e1600377 |
| Geological events | - | - | - |
| Climate change and severe weather | Past and future climate data | Various datasets from the Data Distribution Centre of the Intergovernmental Panel on Climate Change (IPCC) | https://www.ipcc-data.org/index.html |

**Supplementary Information 3.** Methods and data sources for mapping Anthropocene refugia for the American bison and the tiger.

### 1. Species distribution data

a) Current and potential distributions

We retrieved spatial data on current distribution of the American bison (*Bison bison*) and tiger (*Panthera tigris*) from the International Union for Conservation (IUCN) Red List [2]. Information about how these maps were produced can be found in the assessments for each species [8,9]. To represent estimates of where bison and tigers could live today in the complete absence of human influence through time, we used counterfactual estimates of present natural ranges from the recently assembled PHYLACINE database, a phylogeny of all late-Quaternary mammals with extensive functional data and present natural range maps [3]. These maps represent estimates of where species would live today without anthropogenic pressures. Detailed descriptions of how present natural ranges were estimated can be found in the supplemental material of Faurby and Svenning [10], where these ranges were first published.

b) Predicted distribution in 2050

We used the Bioclim predictive algorithm [11] to project future distributions of bison and tiger for 2050 (average 2041–2060). We overcame biases introduced by anthropogenic pressures by generating presence locations at a 20 km separation within present natural range maps from the PHYLACINE v1.2. database [3]. We excluded presence locations for the bison above 4000 m, which is roughly the 95% percentile for upper elevations of mammals reported in the IUCN Red List, and for the tiger above 4500 m, which is the highest elevation for tigers reported in the IUCN. Elevation was determined for the presence locations from the EarthEnv-DEM90 digital elevation model [12].

As potential predictor variables to model the future climatic niche of bison and tiger, we used four bioclimatic variables from the WorldClim dataset at 2.5 arc-minute resolution (www.worldclim.org; [13]). These four variables were maximum temperature of the warmest month, minimum temperature of the coldest month, precipitation of the wettest quarter, and precipitation of the driest quarter. The bioclimatic variables are ecologically meaningful as they represent extremes of temperature and precipitation that are likely to be limiting factors for large mammals' distributions. For future scenarios, we generated a consensus projection using seven general circulation models from the CMIP5 database for RCP8.5 (business as usual). Specifically, the GCMS were BCC-CSM1-1, CCSM4, GFDL-CM3, HadGEM2-ES, IPSL-CM5A-LR, MPI-ESM-LR and NorESM1-1. These GCMs cover a range of CMIP5 model performances and reflect predictions that range from pessimistic to optimistic (IPCC 2013). To threshold potential distributions from Bioclim, we chose a training threshold rule that maximizes the sum of specificity and sensitivity. The performance of Bioclim models was measured using fivefold k-cross-validation to calculate the area-under-the-curve (AUC). AUC values range from ≤ 0.5 for models with discrimination no better than random to 1 for models with perfect discrimination between occupied and unoccupied places [14]. We also inspected threshold-dependent (omission rate for testing points using a threshold set by the 10% training omission rate, OR10) to determine whether models were ovefit, where values close to 0 are not overfit to 1 for models that are overfit [15]. Bioclim predictive performance was high for the bison with an AUC of 0.852, but poor for the tiger with an AUC of 0.562. The OR10 values from the Bioclim models suggested for both bison and tiger that they were not overfit (0.031 and 0.027, respectively.) We restricted our prediction to the area that would potentially be accessible to the species via dispersion by applying a buffer around the present-natural range of 10 degrees (about 1,000km at the equator). This ensures that areas that would naturally be inaccessible to the species (e.g. Africa for tigers) are excluded from the predicted distribution map.

### 2. Anthropogenic pressures

#### a) Current threats

We identified ongoing anthropogenic pressures impacting the American bison and the tiger by using the classification of threats in [7] and the IUCN Red List assessment for each species [8,9] (Table 1). We identified five threat layers for each species that could be used as proxies to map these pressures. For the tiger, war zones and problematic native species and diseases are identified as threats but due to the absence of spatially-explicit data and difficulty to quantify their estimated impact on the species, we didn’t include these threats in the analysis. Information on dams is available from Open Street Map (https://www.openstreetmap.org), but the resolution of our study is too small to identify such localized features, and this threat was thus excluded. Similarly for the American bison, the effect of invasive and other problematic species and as well as storms and flooding are complex to quantify and these threats were not included in the mapping exercise. As new research becomes available detailing in more details the problems that these species face, this list can be updated and the analysis further refined.

We used spatially explicit information on the distribution of anthropogenic pressures from the Human Footprint [16] to map the distribution of threats at the continental level for each species. The Human Footprint maps are globally standardised maps of cumulative human pressures on the natural environment at 1 $km^2$ resolution, available for 8 type of threats (built environments, population density, electric infrastructure, crop lands, pasture lands, roads, railways, and navigable waterways). We chose to apply this methodology for these case studies because it has been found to exhibit an excellent degree of accuracy, especially at identifying threat-free areas [17]. Each pressure layer is scaled based on its estimated impact on the environment (see Table 2 in Venter et al. [16] for more detail on the scaling). For each species, we summed for each pixel the value of relevant pressures and rescaled the resulting score to obtain a threat index between 0 and 1, 1 indicating maximum pressure on the environment.

**Table 1.** Class of threats associated to the American bison (*Bison bison*) and the tiger (*Panthera tigris*), as listed by the IUCN Red List for the two species, and the type of proxy used from the Human Footprint maps.

| Species | Threat | Classification | Proxy used |
|---|---|---|---|
| **American bison** | Agriculture & aquaculture | Annual & perennial non-timber crops | Croplands |
| | | Livestock farming & ranching | Pasture |
| | Transportation & service corridors | Roads & railroads | Roads, Railways |
| | Biological resource use | Hunting & trapping ; Human intrusion & disturbances | Population density |
| | Natural system modification | Other ecosystem modifications | Population density |
| | Invasive and other problematic species, genes and diseases | Invasive non-native/alien species/diseases | NA |
| | | Problematic native species/diseases | NA |
| | Climate change & severe weather | Storms & flooding | NA |
| **Tiger** | Residential & commercial development | Commercial & industrial areas | Built environments |
| | | Tourism & recreation areas | Population density |
| | Agriculture & aquaculture | Annual & perennial non-timber crops | Croplands |
| | | Wood & pulp plantations | Croplands |
| | | Livestock farming & ranching | Pasture |
| | | Marine & freshwater aquaculture | NA |
| | Energy production & mining | Mining & quarrying | NA |
| | Biological resource use | Hunting & trapping terrestrial animals | Population density |
| | Human intrusion & disturbances | Recreational activities | Population density |
| | | War, civil unrest & military exercises | NA |
| | Natural system modifications | Fire & fire suppression | NA |
| | | Dams & water management/use | NA |
| | Invasive and other problematic species, genes and diseases | Problematic native species/diseases | NA |

#### b) Future threats

Spatially-explicit information on projected future anthropogenic pressures were not available for each threat class. However, we were able to map the areas with high probability to be converted into cropland or built areas, using forecasts of agricultural and urban expansion from Oakleaf et al. [18]. In this study, future resource development potentials were created from publicly available global datasets and relatively ranked based on estimated future area expansion derived from past trends [19,20]. This resulted in spatially-explicit information on threat scores based on 1) estimates of fractional amount of agricultural expansion by 2030 extrapolated from 2000–2011 cropland and pasture time series maps and 2) mean probabilities of global urban expansion by 2030, after excluding current urban areas [18]. We used this information to map areas that have a threat score > 50%, i.e. areas that have probability of being converted into cropland or built areas in the future.

### 3. Mapping Anthropocene refugia

We used a threshold to reclassify the continuous map of threat index within the future potential range of the species into a binary map of realised and potential Anthropocene refugia. The threshold corresponds to a certain quantile of the frequency distribution of threat index within the current (IUCN) range of the species. The appropriate threshold to be used is case-specific and should be adapted according to the objectives of the study and the organism considered. A sensitivity analysis showing how results can vary when different thresholds values are used can be useful to identify areas that are consistently predicted as Anthropocene refugia. Here, we report results for a 0.95 quantile in the main text (Box 1, figures D and H), while Figure S1 below shows the results for different threshold values, from the median value (quantile = 0.50) to the maximum value of threat found within the taxa's the current range (quantile = 1). The higher quantile value, the higher value of threat index under which an area is classified as suitable, and the larger the areas identified as Anthropocene refugia. A low threshold will thus provide more conservative maps of Anthropocene refugia (i.e. the value of anthropogenic threat that the species is supposedly able to withstand will be smaller and thus the area identified as refugia will be reduced).

The spatial data and R code used to produce the maps in Box 1 of the main text and Figure S1 of Supplementary Information are available on FigShare (https://doi.org/10.6084/m9.figshare.7845236.v1).

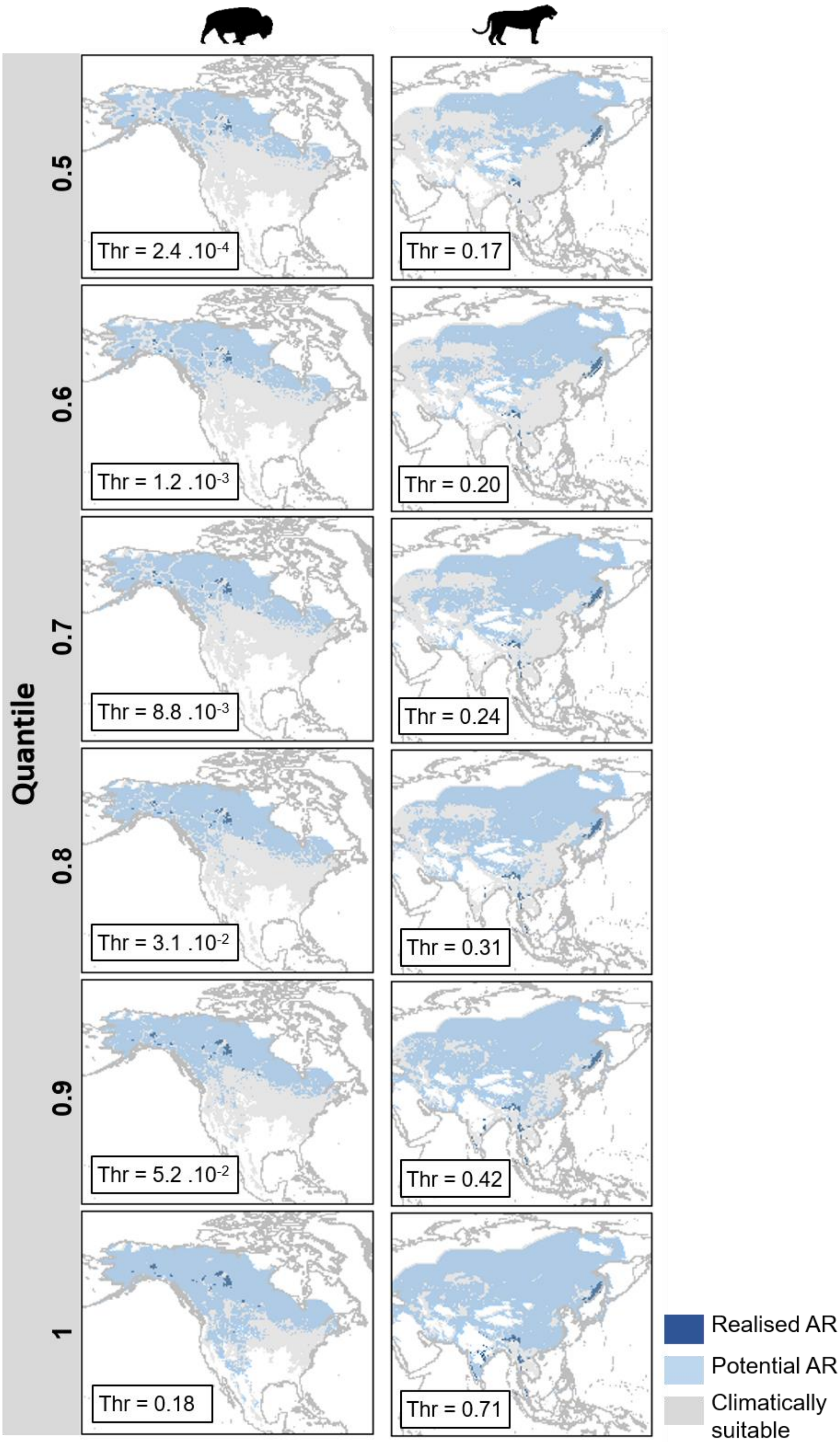


**Figure S1. Maps of realised and potential Anthropocene refugia (AR) for different thresholds of threat index (Thr) for the American bison *Bison bison* and the tiger *Panthera tigris*.** The thresholds used to build the maps correspond to different quantiles of the frequency distribution of threat index within the current range of the species. Realised refugia are indicated in dark blue and potential refugia in light blue. Grey areas are predicted to be climatically suitable for the species in the future but have a value of threat index above the reported threshold.

**REFERENCES**


1. GBIF: The Global Biodiversity Information Facility. 2019 What is GBIF?

2. IUCN. 2018 The IUCN Red List of Threatened Species. Version 2018-2. http://www.iucnredlist.org.

3. Faurby S, Davis M, Pedersen RØ, Schowanek SD, Antonelli A, Svenning J-C. 2018 PHYLACINE 1.2: The Phylogenetic Atlas of Mammal Macroecology. *Ecology* (doi:10.1002/ecy.2443)

4. Enquist BJ, Condit R, Peet RK, Schildhauer M, Thiers BM. 2016 Cyberinfrastructure for an integrated botanical information network to investigate the ecological impacts of global climate change on plant biodiversity. *PeerJ Prepr.* **4**, e2615v2. (doi:10.7287/peerj.preprints.2615v2)

5. Maitner BS *et al.* 2018 The BIEN R package: A tool to access the Botanical Information and Ecology Network (BIEN) database. *Methods Ecol. Evol.* **9**, 373–379. (doi:10.1111/2041-210X.12861)

6. Kaschner K, Kesner-Reyes K, Garilao C, Rius-Barile J, Reest T, Froese R. 2016 AquaMaps: Predicted range maps for aquatic species. *World Wide Web Electron. Publ.* **Version 08/2016**.

7. Salafsky N *et al.* 2008 A Standard Lexicon for Biodiversity Conservation: Unified Classifications of Threats and Actions: *Classifications of Threats & Actions*. *Conserv. Biol.* **22**, 897–911. (doi:10.1111/j.1523-1739.2008.00937.x)

8. Aune K, Jørgensen D, Gates C. 2017 *Bison bison* (errata version published in 2018). The IUCN Red List of Threatened Species 2017: e.T2815A123789863. (doi:http://dx.doi.org/10.2305/IUCN.UK.2017-3.RLTS.T2815A45156541.en)

9. Goodrich J *et al.* 2015 *Panthera tigris*. The IUCN Red List of Threatened Species 2015: e. T15955A50659951. (doi:http://dx.doi.org/10.2305/IUCN.UK.2015-2.RLTS.T15955A50659951.en)

10. Faurby S, Svenning J-C. 2015 Historic and prehistoric human-driven extinctions have reshaped global mammal diversity patterns. *Divers. Distrib.* **21**, 1155–1166. (doi:10.1111/ddi.12369)

11. Nix H, Busby J. 1986 BIOCLIM, a bioclimatic analysis and prediction system. *Div. Water Land Resour. Canberra*

12. Robinson N, Regetz J, Guralnick RP. 2014 EarthEnv-DEM90: A nearly-global, void-free, multi-scale smoothed, 90m digital elevation model from fused ASTER and SRTM data. *ISPRS J. Photogramm. Remote Sens.* **87**, 57–67.

13. Hijmans RJ, Cameron SE, Parra JL, Jones PG, Jarvis A. 2005 Very high resolution interpolated climate surfaces for global land areas. *Int. J. Climatol.* **25**, 1965–1978. (doi:10.1002/joc.1276)

14. Fielding AH, Bell JF. 1997 A review of methods for the assessment of prediction errors in conservation presence/absence models. *Environ. Conserv.* **24**, 38–49.

15. Anderson RP, Gonzalez I. 2011 Species-specific tuning increases robustness to sampling bias in models of species distributions: An implementation with Maxent. *Ecol. Model.* **222**, 2796–2811. (doi:10.1016/j.ecolmodel.2011.04.011)

16. Venter O *et al.* 2016 Global terrestrial Human Footprint maps for 1993 and 2009. *Sci. Data* **3**, 160067. (doi:10.1038/sdata.2016.67)

17. Allan JR, Venter O, Watson JEM. 2017 Temporally inter-comparable maps of terrestrial wilderness and the Last of the Wild. *Sci. Data* **4**, 170187. (doi:10.1038/sdata.2017.187)

18. Oakleaf JR, Kennedy CM, Baruch-Mordo S, West PC, Gerber JS, Jarvis L, Kiesecker J. 2015 A World at Risk: Aggregating Development Trends to Forecast Global Habitat Conversion. *PLOS ONE* **10**, e0138334. (doi:10.1371/journal.pone.0138334)

19. Seto K, Güneralp B, Hutyra LR. 2015 Global Grid of Probabilities of Urban Expansion to 2030.

20. Ramankutty N, Foley JA. 1999 Estimating historical changes in global land cover: Croplands from 1700 to 1992. *Glob. Biogeochem. Cycles* **13**, 997–1027. (doi:10.1029/1999GB900046)
